# Photoactivation of olfactory sensory neurons does not affect action potential conduction in individual trigeminal sensory axons innervating the rodent nasal cavity

**DOI:** 10.1101/517391

**Authors:** Margot Maurer, Nunzia Papotto, Julika Sertel-Nakajima, Markus Schueler, Frank Möhrlen, Karl Messlinger, Roberto De Col, Stephan Frings, Richard W Carr

## Abstract

Olfactory and trigeminal chemosensory systems reside in parallel within the mammalian nose. Psychophysical studies in people indicate that these two systems interact at a perceptual level. Trigeminal sensations of pungency mask odour perception, while olfactory stimuli can influence trigeminal signal processing tasks such as odour localization. While imaging studies indicate overlap in limbic and cortical somatosensory areas activated by nasal trigeminal and olfactory stimuli, there is also potential cross-talk at the level of the olfactory epithelium, the olfactory bulb and trigeminal brainstem. Here we focused on potential interactions between olfactory and trigeminal signaling in the nasal cavity. We first used a forced choice paradigm to ascertain whether trigeminal and olfactory stimuli could influence behavior in mice. Mice avoided water sources associated with volatile TRPV1 and TRPA1 irritants, however, this aversion was mitigated when combined with a pure odorant (rose fragrance, phenylethyl alcohol, PEA). To determine whether olfactory-trigeminal interactions within the nose could potentially account for this behavioral effect we recorded from single trigeminal sensory axons innervating the nasal epithelium using an isolated in vitro preparation. To circumvent non-specific effects of chemical stimuli, optical stimulation was used to excite olfactory sensory neurons in a mouse line expressing channel-rhodopsin under the olfactory marker protein. During photoactivation of olfactory sensory neurons there was no modulation of action potential conduction in individual trigeminal axons. Similarly, no evidence was found for trigeminal axon collateral branching that might serve as a conduit for cross-talk between the olfactory epithelium and olfactory dura mater. Using direct assessment of trigeminal signals emanating from the mouse nasal cavity we see no evidence for paracrine nor axon reflex mediated cross-talk between olfactory and trigeminal sensory systems in the nasal cavity.

## Introduction

The sensory innervation of the mammalian nasal cavity by the trigeminal and the olfactory systems endows the nasal epithelia with a broad spectrum of sensory modalities. Trigeminal fibers originating from the ethmoid and nasopalatine nerves [1] detect irritants, temperature changes and mechanical stimuli [2, 3], while olfactory receptor neurons respond specifically to odorants and non-specifically to mechanical stimuli [4]. In addition to the extended trigeminal network innervating the nasal respiratory epithelium, the olfactory sensory epithelium also contains trigeminal fibers [5-8] [9]. In co-ordination with the olfactory system, trigeminal chemesthesis contributes to a continual analysis of the composition of the inhaled air for harmful and useful compounds with the trigeminal signaling being implicated in the induction of protective reflexes [10], pain perception [11] and subsequent behavioural responses.

It has been known for a long time that nasal olfactory and trigeminal sensory systems interact with one another on multiple levels of information processing, beginning with the fact that most odorants can stimulate trigeminal fibers and that most irritants have an odour [12]. Work on human nasal sensation has led to the concept that chemical stimulation of the nose triggers a multimodal response, which is more appropriately described as an integrated afferent signal rather than as two separate streams of trigeminal and olfactory information [12-22]. Most studies have focused on the suppressive effect of trigeminal stimuli that elicit sensations of pungency, on the perception of odorants. Similarly, in animal models, the impact of trigeminal activation on olfactory signaling has been examined and release of calcitonin gene-related peptide (CGRP) from trigeminal afferents can suppress excitatory signals in olfactory sensory neurons [5, 23]. However, it is also possible that olfactory signaling can influence signaling in trigeminal sensory neurons [24, 25]. For example, the sensory task of spatial localization of odours attributed to activation of trigeminal neurons is enhanced by odorants [26]. Presently, there is no clear understanding of the molecular pathways nor the anatomical sites at which olfactory signaling might affect trigeminal activity. Imaging studies in people indicate overlap of central trigeminal and olfactory processing pathways [27]. Clinical evidence indicates that olfactory stimuli can affect the course of primary headache disorders, in particular in migraine [28-33] for which modulation within the trigeminal brainstem nuclei is implicated [33, 34].

Here we systematically explored the effect of olfactory stimulation on trigeminal signaling in the nose. We used anatomical and electrophysiological techniques to characterize the ethmoid nerve innervation of the nasal cavity at a single fibre level. To examine directly the influence of olfactory stimuli on trigeminal signaling we used an optogenetic mouse line expressing channel rhodopsin under the olfactory marker protein promoter (OMP-ChR2- YFP) that allowed selective photoactivation of olfactory sensory neurons. Using photoactivation of olfactory sensory neurons we were unable to detect any effect on action potential signaling in single trigeminal sensory afferent from the nasal cavity and conclude that olfactory sensory neurons exert minimal influence on trigeminal signals within the nasal cavity.

## Methods

Animal housing and all experimental procedures were carried out in compliance with the guidelines for the welfare of experimental animals as stipulated by the Federal Republic of Germany. For behavioural experiments ethical approval was obtained under approval number 35-9185.81/G-104/16.

### Animals

Transgenic mouse lines were provided by Thomas Bozza (Northwestern University, Evanston, IL, USA) for OMP-ChR2-YFP mice [35], by David D. McKemy (University of Southern California, San Diego, CA, USA) for TRPM8-eGFP mice [36] and by Rohini Kuner (University Heidelberg, Heidelberg, Germany) for SCN10A- Cre::Rosa26-tdTomato mice [37]. Exclusively male C57BL/6N (Charles River Laboratories) mice were used for behavioral experiments.

### Ex vivo nasal cavity preparation

Adult C57BL/6N mice of both sexes and with body weights ranging from 22 to 31 g were anaesthetized with isoflurane (Abbott, Weisbaden, Germany) in a sealed glass chamber (2 litre volume) and subsequently killed by cervical dislocation. The head and lower jaw were removed and the cranial vault cleared of overlying skin and muscle. Similar to our previous description of an ex vivo half skull preparation for recordings from meningeal afferents [38], the skull was divided in the sagittal plane with a scalpel. The cortex, brain stem and olfactory bulb were removed along with the nasal septum. Each half skull was embedded in a perspex chamber using agarose (8%; Sigma, Munch, Germany) such that the nasal cavity and the bony cavity of the olfactory bulb formed a contiguous tissue bath. The average experimental recording time for each half skull was 2-4 hours.

The half skull bath was perfused continuously at ca. 4 ml.min^−1^ with physiological solution comprising (in mM): hydroxyethyl piperazine ethanesulfonic acid solution (HEPES), 6; NaCl, 118; KCl, 3.2; NaGluconate, 20; D-Glucose 5.6; CaCl2, 1.5; MgCl2, 1. The pH was adjusted to 7.4 with NaOH. The temperature of the perfusing solution was controlled at 32.0 ± 0.5 °C with an in-line resistive heating element regulated by feedback from a thermocouple positioned in the bath.

### Recording arrangement

The anterior ethmoid nerve was identified along its course within the dura mater of the anterior cranial fossa which it enters proximally through the anterior ethmoid foramen and exits through the cribroethmoid foramen to enter the nasal cavity [39].

The nerve was cut as close to its exit through the ethmoid foramen as possible and the distal cut end freed of surrounding dura over a length of approximately 4 mm sufficient to attach a glass recording electrode to the cut end by light suction. The glass recording electrode was filled with physiological solution and the tip cut with a sapphire blade to match the diameter of the ethmoid nerve. Signals were recorded over the sealing resistance relative to a Ag/AgCl pellet in the bath using a differential amplifier (NL104A, Digitimer, City, UK). Signals were filtered (low-pass 5 kHz, 80 dB Bessel), digitized (20 kHz, micro 1401, Cambridge Electronic Design, Cambridge, UK) and stored to disk for subsequent analysis.

### Mechanical, electrical, thermal and chemical stimulation

Receptive fields of individual sensory axons in the nasal cavity were either established broadly using a mechanical (von Frey) stimulus or in some cases using an electrical stimulator without prior mechanical searching. In the case of mechanical searching, a servo driven mechanical stimulator [40] was placed at sites within the area mapped with the von Frey filament. The mechanical stimulator was used to deliver brief sinusoidal (10ms pulse width) mechanical stimuli at different sites until a single unit response was identified. For electrical stimulation a rayon insulated platinum iridium wire (ISAOHM, Isabellenhütte, Dillenburg, Germany), 20 µm in diameter and exerting a buckling load of ca. 0.4 mN, was placed on the tissue and served as a the cathode. A Ag/AgCl pellet (WPI, Sarasota, Florida, USA) positioned in the tissue bath served as the anode for constant current electrical stimuli (1 ms, <100µA). Thermal stimuli were delivered by changing the temperature of the solution perfusing the bath. The heating-element bath perfusion circuit had a thermal time constant of approximately 14s. For chemical stimuli, substances were delivered to the solution perfusing the bath, excluding ammonia (NH_3_).

Ammonia (NH_3_) was applied to the nasal cavity in volatile form. For this series of experiments the half skull was mounted in the recording bath slightly inclined in the sagittal plane, such that the fluid level in the bath could be reduced transiently to expose most of the nasal cavity to air leaving sufficient solution to maintain fluid around the electrophysiological recording pipette attached to the ethmoid nerve within the anterior cranial fossa. Ammonia was then applied by brief pressure pulses to a 2 ml syringe approximately one quarter filled with ammonium (NH_4_Cl) solution (4.3% w/v) to deliver ammonia vapour.

### Determination of axonal conduction velocity

Axonal conduction velocity was calculated by dividing the latency of the action potential response to electrical stimulation by the length of axon between the stimulating and recording sites. The length of nerve between the two sites was estimated visually by reference to a graticule placed in the light path of the microscope’s ocular objective.

### Determination of mechanical threshold

Estimates of mechanical activation threshold were determined for individual axons by determining the likelihood of an action potential response at several discrete stimulus strengths as previously described [40]. Briefly, the probability (number of responses/number of stimuli) of evoking an action potential response across five repeat presentations at each force was determined and the regression of probability on force was fit with a sigmoid function. Mechanical threshold was taken as the inflection point of the fit. Mechanical stimuli were sinusoidal in form and typically of 10 ms duration. The force of mechanical stimulation was taken as the peak maximum of the sinusoidal force profile and force was divided by tip area (200 µM diameter, 0.125 mm^2^) to estimate mechanical stress.

### Evaluation of response to temperature and chemical stimuli

Extracellular recordings of single C-fibres from peripheral nerves are typically performed by manually teasing the cut end of a nerve into progressively smaller filaments. However, short length of nerve and limited access preclude use of split fibre techniques for the ethmoid nerve in the mouse. We therefore adopted a loose extracellular patch technique to record from the entire ethmoid nerve. In this configuration, signals from multiple units were recorded. To refine this to a single fibre recording a small electrical or mechanical stimulus is delivered to the tissue (olfactory or respiratory epithelium in this case) until we established a time-locked single fibre response from stimuli applied within the receptive field. Even with a time-locked action potential response, the small amplitude and relative uniform shape of extracellularly recorded C-fibre action potentials make it difficult to discern firing patterns of individual axons. Therefore, to ascertain whether functionally identified single axons responded to thermal or chemical stimuli we used the technique of latency “marking” that relies on an increase in axonal conduction velocity in C-fibres subsequent to each action potential [41]. To utilise this physiological principle, constant frequency electrical stimulation was delivered to the receptive field of an individual axon before and during application of thermal or chemical stimuli. Units were considered to have responded with the generation of action potentials if the latency of response to electrical stimulation increased or if the axon became transiently refractory to electrical stimulation, i.e. no response to electrical stimulation was evident.

### Search for axon collaterals

Previous studies using peripheral dye injections into the olfactory bulb and nasal cavity resulted in double-labelled neurons in the rat trigeminal ganglion and suggested that individual trigeminal axons branch divergently to innervate both the olfactory epithelium and the olfactory bulb [42]. Consequently, we used functional techniques to examine the extent of divergent axonal branching in the ethmoid nerve, which was assumed to innervate also the dura mater of the anterior cranial fossa surrounding the olfactory bulb dorso-laterally and termed here “olfactory dura mater”. The first paradigm used the same preparation for recording single afferents from the distal cut end of the ethmoid nerve as described above. Time-locked electrically-evoked action potentials were used to identify receptive fields within either the olfactory epithelium or the olfactory dura mater. In addition, the nasal cavity or the olfactory dura mater was stimulated mechanically with a von Frey hair. Mechanical stimuli were applied over the full spatial extent of the anterior cranial fossa by slowly probing sequential sites.

In a second series of experiments, functional verification of axon collaterals was sought by changing the site of recording to the proximal cut end of the anterior ethmoid nerve at a site immediately distal to its traverse of the cribriform plate through the cribroethmoid foramen. In this configuration, any action potentials recorded in the anterior ethmoid nerve in response to mechanical stimulation of the olfactory dura mater must be travelling anti-dromically via axon reflex between branches of individual axons.

### Behavioural assessment of volatile chemical stimuli on water consumption

In order to avoid confounding influences of hormonal changes on olfaction, exclusively male C57BL/6N mice aged from 9-14 weeks were used for behavioral testing [43, 44]. The housing facility was kept at a constant temperature of 22 ± 1 °C with a 12 h light/dark cycle. To assess the influence of irritant and odorant stimuli a forced choice paradigm was developed. Two water bottles were positioned inside home cages separated by a distance of 8 cm and in all respects indistinguishable (height above cage floor, resistance to water flow, hydrostatic pressure determined by starting water volume, nozzle diameter and orientation in cage).

Prior to testing, mice were acclimatized for seven consecutive days in individual ventilated polycarbonate cages (32 cm × 16 cm × 30 cm, L × W × D) with access to water from two bottles and food *ab libitum*. Volatile odorant and irritant stimuli were added to the drinking tubes on day eight. Felt washers were soaked in the volatile agents and the washers were placed in perforated aluminum containers with a central hole such that the housing rings could be pushed onto the sipping tubes of water bottles (Figure 1A). This system exposed the immediate vicinity of the drinking tube to a high concentration of volatile compound. Odorant and irritant stimuli remained around the sipping tubes for a period of 24 hours (Figure 1B). Water bottles were washed daily with ethanol and water and refilled with fresh water before being replaced in the cage with a felt washer freshly soaked in odorant or irritant. The amount of water consumed was quantified by establishing the change in weight of each bottle over each consecutive 24 hour period during acclimatization, exposure to volatile agents and in the post-exposure phase. Relative values of water consumption were determined as: water consumption test bottle / (consumption test bottle + consumption control bottle).

**Figure 1:**
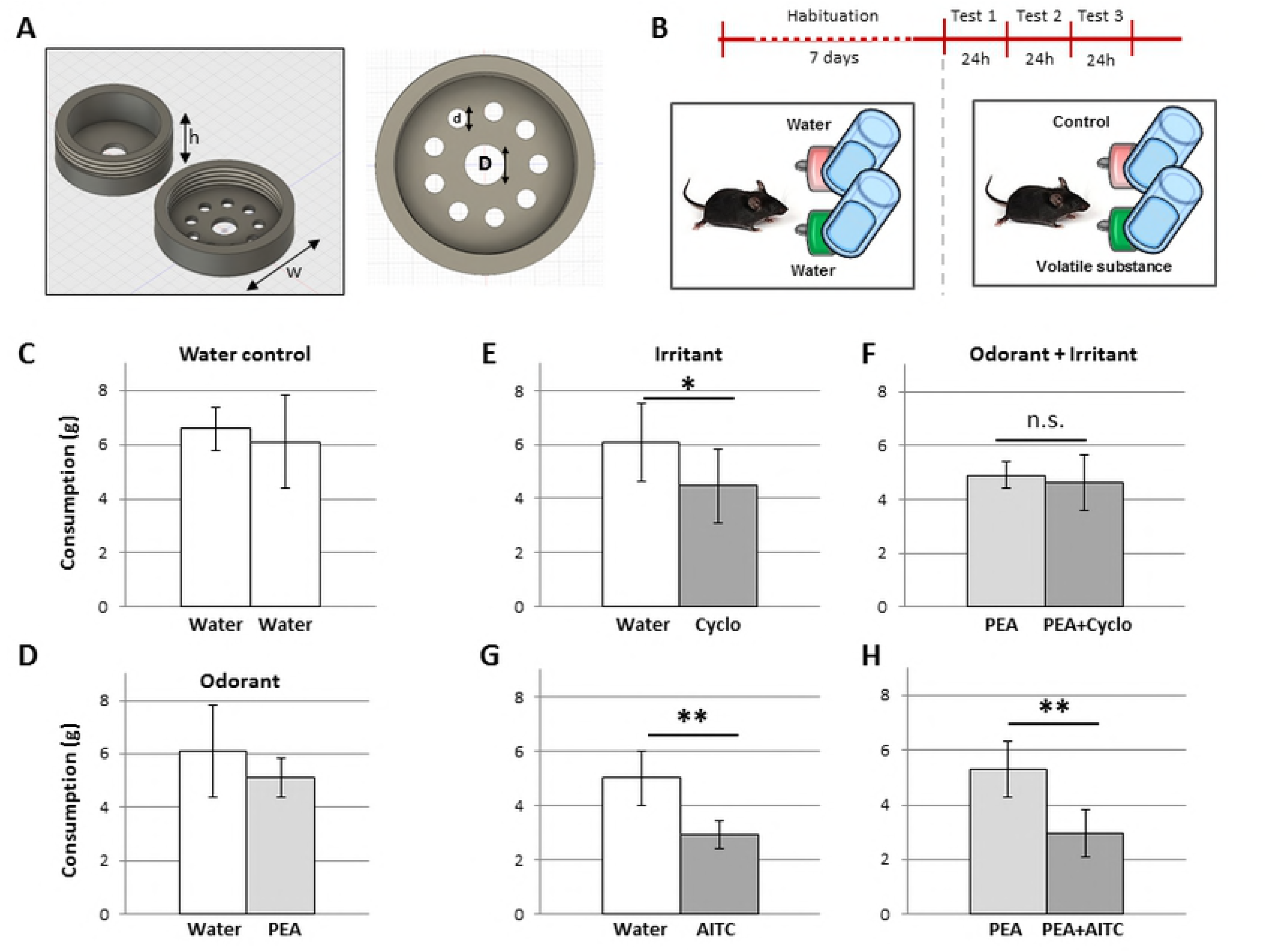
Water preference paradigm to assess olfactory-trigeminal interactions using volatile irritants and odorants in freely moving mice. Annular pieces of felt were soaked in irritant and odorant solutions and placed in an aluminum housing (A) that was mounted around each of two water sipping tubes made available to mice in their home cages (B). The dimensions of the annular housing indicated in panel A are h= 11mm; w= 30mm; D= 8.5mm and d= 4mm. Mice drank *ad libitum* from either of two water sources throughout the experiment. Following a seven day acclimatization period (B, upper) irritant and/or odorant was added to the aluminum casings around one or both water sources for three consecutive days in a randomized (left/right) fashion (B). In the control condition for two sources of water alone, there was no difference in the amount of water consumed from each bottle (C). Similarly, there was no difference in water consumption between water and water combined with the pure odorant PEA (D). Mice consumed less water from sources of water surrounded by an irritant, either cyclohexanone (E) or AITC (G). In combination with the odorant PEA mice consumed less from a source surrounded by AITC (H) but did not avoid a source surrounded by cyclohexanone (F).

### Anterograde tracing

The technique of anterograde tracing has been described previously [45]. Briefly, rats were killed by inhaling CO_2_, the head removed, the cranial vault cleared of overlying tissue and subsequently hemisected mid-sagittally. The brain was removed sparing the cranial dura mater and trigeminal ganglion. The ethmoid nerve was dissected free about 2 mm beyond its traverse into the anterior cranial fossa through the ethmoid foramen (Figure 2A). A small crystal of the carbocyanine dye Di-I3 (1,1′Dioctadecyl-3,3,3′,3′-Tetramethylindocarbocyanine Perchlorate, D282, Molecular Probes, Eugene, OR, USA) was placed on the distal cut end of the ethmoid nerve and covered with a piece of gelatin sponge (Abgel, Sri Gopal Labs, Mumbai, India) to avoid spreading of the dye. Dye was transported at approximately 2.5mm/week and to trace the length of anterior ethmoid nerve took around 2 months.

### Immunohistochemistry

Mice heads were prepared for immunohistochemistry using the same procedure as that used for electrophysiological experiments (see above). The cranial vault was fixed in 4% paraformaldehyde for 2 hours. The olfactory epithelium was isolated using the deboning protocol described in Dunston, Ashby (46). Trigeminal ganglia and the ethmoid nerve were isolated from half-skulls with the cortices, brainstem and olfactory bulb removed.

Samples were dehydrated in a 10% sucrose solution (10% sucrose (w/v), 0.05% NaN_3_ (w/v) in PBS, pH=7.4) for 2 hours and cryoprotected in 30% sucrose solution (30% saccharose (w/v), 0.05% NaN_3_ (w/v) in PBS, pH=7.4) overnight before being embedded in Optimal Cutting Temperature medium (OCT, Sakura Finetek, CA, USA) and stored at −20°C before sectioning. Tissue was sectioned serially at 25 µm on a cryotome (Thermo Scientific Microm HM 550, Germany). Sections were mounted onto glass slides (Superfrost Plus™, Thermo Fisher Scientific). Whole mounts were washed in 0.1 M phosphate buffer and blocked with 5% ChemiBLOCKER (EMD Millipore Billerica) containing 0.5% Triton X-100 and 0.05% NaN 3 for 1 h. Primary antibodies were applied overnight at 4 °C.” Sections were washed in 0.1M PBS and blocked with 5% Chemiblocker (Millipore, Darmstadt,Germany) comprising 0.5% Triton X-100 and 0.05% NaN3 in PBS at pH=7.4. Primary antibodies (Table 1) were applied overnight in 5% Chemiblocker containing 0.5% Triton X-100 and 0.05% NaN_3_ in PBS (pH=7.4) For fluorescent labeling section were incubated with secondary antibodies: Alexa Fluor 488-labelled goat anti-rabbit (dilution 1:1000, Invitrogen) and 1μg/ml DAPI (DAPI, dilactate ≥98%, Sigma-Aldrich, Germany). Slides were mounted with fluorescence mounting medium (Dako, Agilent, Italy) and imaged (Nikon Eclipse 90i/C1, Nikon, Japan) and analyzed (NIS elements, version 4.0; Nikon) using confocal techniques.

**Table 1:** Primary antibodies

| ***Antibody*** | ***Host*** | ***Supplier*** | ***Catalogue number*** | ***dilution*** |
| --- | --- | --- | --- | --- |
| Anti-GFP | Rabbit (polyclonal) | Abcam<br>(Cambridge, UK) | Ab6556 | 1:400 |
| Anti-RFP | Rabbit (polyclonal) | Rockland<br>(Limerick, USA) | 600-401-379 | 1:200 |
| Anti-TRPA1 | Rabbit (polyclonal) | Abcam<br>(Cambridge, UK) | ab58844 | 1:400 |

### Chemicals

L-menthol, allyl isothiocyanate (AITC), Capsaicin, cyclohexanone, phenylethyl alcohol (PEA, 2-phenylethylethanol) and ammonium chloride (NH_4_Cl) were all purchased from Sigma-Aldrich, Munich, Germany. Substances were made up in stock solutions of either DMSO, alcohol or mineral oil. Stock solutions were diluted to the required concentration in the perfusing solution on the day of each experiment. Ammonium chloride was made up as a 4.3% (w/v) solution in distilled water and applied as ammonia (NH_3_) vapour.

### Data analysis and statistics

Electrical stimulation protocols were tracked online using custom scripts in Spike2 (CED, Cambridge, UK) and analyzed offline (IgorPro, Lake Oswego, OR, USA).

For statistical comparisons between groups Student’s t-test were used. For multiple group comparisons, 2-way ANOVA was used with post-hoc Tukey HSD for pair-wise comparisons within factors. P values less than 0.05 were deemed significant.

## Results

### Behavioural assessment of olfactory and trigeminal chemosensory interaction

The influence of odorant activation of olfactory sensory neurons on trigeminal signal processing was assessed at a behavioural level in mice using a forced-choice water consumption paradigm. Home cages were outfitted with two identical water bottles for daily water consumption. While the water itself was not contaminated, access to the water source could be influenced by the presence of a volatile agent around the sipping tube. Chemical stimuli were applied to a felt ring that was encased in aluminum housing, annular in form and positioned near the tip of the sipping tube inside the cage (Figure 1A). Mice were acclimatized to the presence of two water bottles with empty aluminum housings over a seven day period before assessment of the preference in presence of an odorant, an irritant or both volatile substances combined (Figure 1B). Using commercially available drinking bottles we were able to establish a baseline condition in which the amount of water consumed from the two water sources did not differ (Figure 1C, paired t-test, n=10, p=0.50). Addition of the odorant PEA to one bottle reduced the overall water consumption in that cage compared with just two water bottles (cf. Figure 1C and 1D; ANOVA, factor time F(1,9)=6.43, p=0.016) but did not affect drinking preference within individual cages (Figure 1D, ANOVA interaction time*PEA F(1,9)=2.12, p=0.154377). In contrast to the odorant PEA, the presence of irritant compounds resulted in an aversion from the affected water bottle. For the TRPA1 agonist AITC, there was an aversion in both the presence (Figure 1H) and absence (Figure 1G) of PEA (ANOVA, F(1,9)=56.70, df=39, p<0.001 factor AITC). For the TRPV1 agonist [47] cyclohexanone there was also an aversion when presented side-by-side with water alone (Figure 1E, ANOVA, F(1,11)=6.99, df=47, p=0.0011 factor cyclo, post-hoc Tukey HSD water vs cyclohexanone p=0.014) but the aversion to cyclohexanone was masked when PEA was added to both water sources (Figure 1F post-hoc Tukey HSD PEA vs PEA+cyclohexanone, p=0.94). However, the statistical interaction between PEA and cyclohexanone did not reach statistical significance (ANOVA, interaction PEA*cyclo F(1,11)=3.44, p=0.07). Nevertheless, this behavioural effect consolidated the proposal that odorant olfactory stimuli can mitigate aversion to volatile irritants in mice. Accordingly, we designed a protocol to establish whether this cross-modal interaction might occur within the nose.

### Nasal and dural projections of the anterior ethmoid nerve

We used the dextran amine DiI as an anterograde tracer [45] to establish initially the meningeal and intra-nasal innervation arising from the ethmoid nerve in the rat (Figure 2A). To further specify ethmoid axons innervating the nasal cavity, we imaged tissue from wildtype mice after immunostaining for TRPA1 (Figure 2C) and from mouse reporter lines for TRPM8 (Trpm8^eGFP^, Figure 2B) and the sodium channel isoform NaV1.8 (Scn10a::tdTomato, Figure 2D). In the trigeminal ganglion (Figure 2C-E, right panels) and the anterior ethmoid nerve (Figure 2C-E, centre panels) we observed respectively somata and axons expressing Trpm8 (Figure 2C), Trpa1 (Figure 2D) and NaV1.8 (Figure 2E). Within the olfactory epithelium (Figure 2C-E, left panels) we observed individual somatosensory nerve terminals positive for Trpm8 (Figure 2C, left) and Trpa1 (Figure 2C, left) that traversed the olfactory epithelium from the lamina propria to the apical surface of the olfactory epithelium as a single unbranched axon. Trpm8 and Trpa1-positive axons were more often encountered in sections from the posterior reaches of the nose.

**Figure 2:**
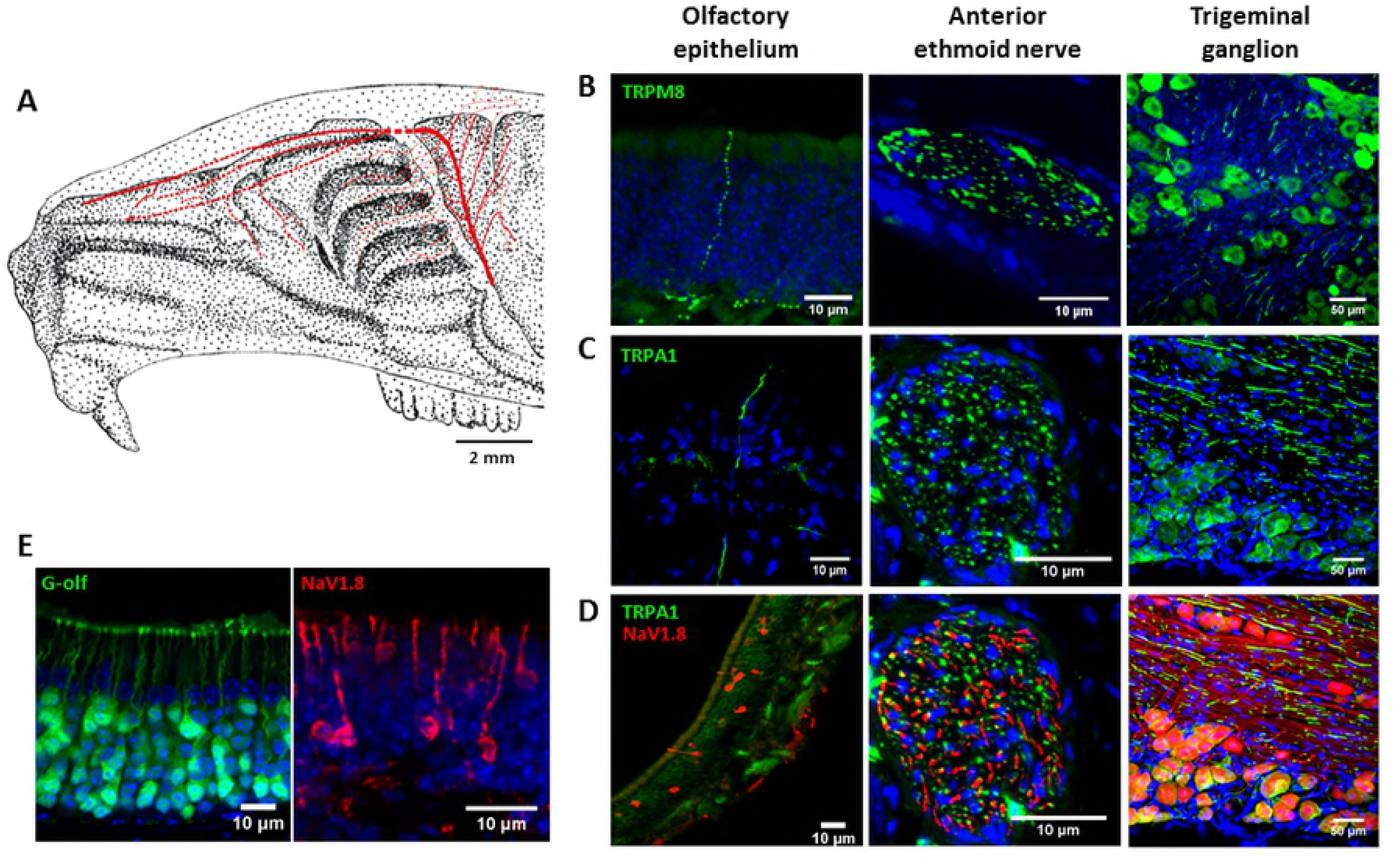
Structural characterization of the ethmoid innervation of the rodent nasal cavity. Distribution of ethmoid nerve branches revealed by anterograde tracing with the dextran amine DiI (A, red tracing) overlaid on a sketch of the rat skull in mid-sagittal section (A). TRPM8-expressing (B, green), TRPA1-immuno-positive (C, green) and NaV1.8-expressing (D, red) structures within the olfactory epithelium (C-E, left panels), the anterior ethmoid nerve (C-E, centre panels) and the trigeminal ganglion (C-E, right panels). NaV1.8-positive structures resembling trigeminal axons were not observed within the olfactory epithelium (E, left). However, within the olfactory epithelium cells expressing tdTomato in SCN10A::tdTomato mice (E, left) were morphologically similar to olfactory sensory neurons expressing the transducer protein G-olf (E, right). Blue in panels B-E are DAPI.

However, although NaV1.8-positive structures were present in sections of olfactory epithelium from Scn10a::tdTomato mice (Figure 2E, left) the labeled cells did not resemble somatosensory axons. Instead, NaV1.8-positive cells within the olfactory epithelium (Figure 2E, left) had morphologies consistent with olfactory sensory neurons as can be seen when compared with OSNs labelled with antibodies to the olfactory G-protein (G-olf, Figure 2E, right). The lack of NaV1.8-positive trigeminal axons within the olfactory epithelium (Figure 2D left and Figure 2E, left) may be influenced by low levels of fluorescent signal in small single axon structures.

### Characterization of the anterior ethmoid innervation of the nasal cavity

We recorded extracellular action potential signals from the distal cut end of the anterior ethmoid nerve (Figure 3A) and identified 71 individual axons with mechanical or electrical receptive fields in the nasal cavity. Consistent with structural reports indicating a preponderance of thinly myelinated and unmyelinated axons in the anterior ethmoid nerve [10] we recorded signals from 8 A-delta (≥1.5m/s) axons and 63 C-fibre axons (<1.5m/s) in the ethmoid nerve with axonal conduction velocities ranging from 0.22 to 2.0m/s (Figure 3B). Thirty-one units were identified using a servo-driven mechano-stimulator. All mechanically-sensitive units had receptive fields within the respiratory epithelium (Figure 3C) and had conduction velocities spanning 0.28-2.0m/s. Absolute mechanical threshold was tested in eight units and ranged from 0.8-8.6mN. Mechanical receptive fields were punctate, contiguous and comprised areas of approximately 0.1-0.8 mm^2^. Of the 31 mechanically sensitive units, 13 were polymodal, with 6 units showing additional heat sensitivity (dark blue markers, Figure 3E) and 11 were also chemosensitive, responding to either capsaicin (250 nM; 6/6), menthol (10 µM; 1/3), allyl isothiocyanate (AITC, 20 µM; 3/4) or ammonia vapour (NH_3_, 5/5). Using an electrical search stimulus, an additional 40 single units were identified with conduction velocities ranging from 0.23-1.11m/s. Sixteen of these units were polymodal; 6/10 and responded to either mechanical stimuli delivered by the electrical probe itself (i.e. 0.4 mN), heating (12/14) or chemical stimuli (9/20), responding to either capsaicin (250 nM; 2/4), menthol (10 µM; 1/2), or cyclohexanone (1% w/v; 4/9) or ammonia vapour (NH_3_, 5/5). We did not record electrically-evoked signals from any cold-sensitive axons. However, background activity consistent with cold-sensitive axons in the ethmoid nerve, i.e. burst activity that was increased upon cooling and silencing during warming, was observed in three preparations (data not shown).

**Figure 3:**
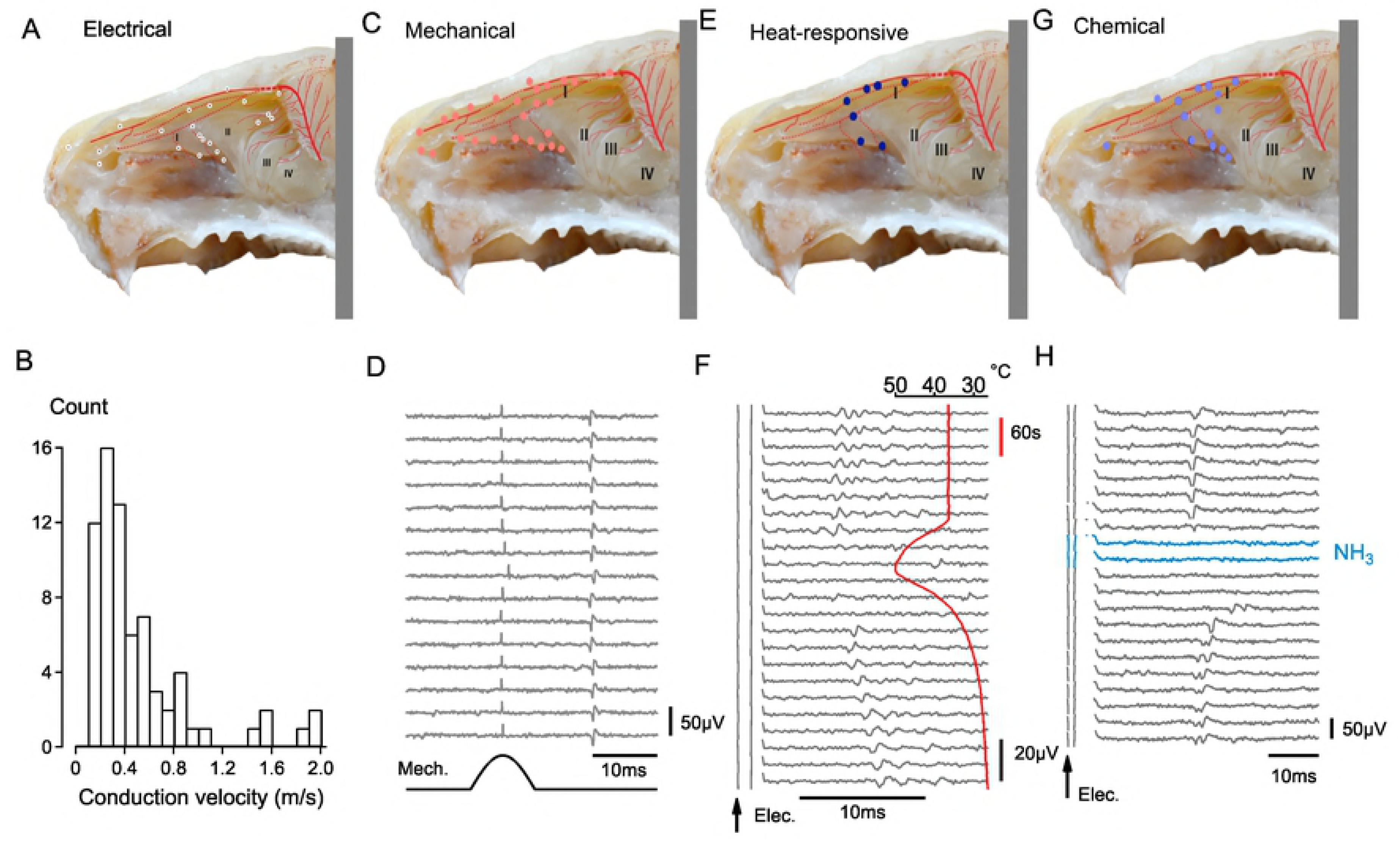
Functional assessment of individual sensory axons in the ethmoid nerve. Extracellular signals were recorded from axons in the ethmoid nerve with receptive fields in the respiratory and olfactory epithelia lining the nasal cavity with axonal conduction velocities ranging from 0.2-2m/s (B). The receptive fields of individual units activated by electrical (A, white markers), mechanical (C, pink markers), heat (E, dark blue markers) and chemical (G, light blue markers) stimuli are shown together with an example response of an individual trigeminal afferent to mechanical stimulation (D), heating (F) and ammonia vapour (H, NH_3_). The red trace in panel F indicates temperature.

### Influence of olfactory sensory neuron photoactivation on single trigeminal sensory afferents within the nasal cavity

To dissociate trigeminal and olfactory chemosensory systems we used photostimulation to activate olfactory sensory neurons (OSN) in isolation and synchronously using preparations from OMP/ChR2-YFP mice (Figure 4A&B)[48]. To establish the efficacy of photostimulation of OSNs we recorded electro-olfactogram (EOG) signals from the surface of the olfactory epithelium during stimulation of the tissue with sinusoidal pulses of blue light (473 nm; Figure 4C). By varying stimulus pulse width we found a peak in the EOG amplitude for pulse widths between 10-20 ms (Figure 4D). In addition, prolonged OSN photoactivation elicited an EOG with an initial phasic component and a sustained tonic component (Figure 4C&D). We thus used photostimulation pulse widths of 10 ms, corresponding to the most synchronous OSN activation (Figure 4D), and 100 ms, corresponding to the approximate length of a sniffing cycle in the mouse [49]. The effect of OSN photoactivation on single trigeminal afferent signals was determined during a 10ms light pulse (Figure 4E), corresponding to the peak amplitude of the photoevoked EOG (Figure 4D) and 100ms, to replicate sustained OSN activation. For seven trigeminal C-fibre axons, the latency of electrically-evoked action potential responses was not altered with repeat applications of a 10-100 ms photoactivation of olfactory sensory neurons (Figure 4E). For comparisons between individual trigeminal afferents we determined the average response to electrical stimulation (grey open markers, Figure 4F) and compared these to the latency of action potentials signals following paired light and electrical stimulation (blue traces, Figure 4E and blue markers, Figure 4F). Taking the ratio of these two latencies (Figure 4G) we saw no effect (paired t-test, n=7, p= 0.29) of OSN photostimulation on trigeminal axonal conduction (Figure 4H). Consistent with this result, we also observed no change in the electrical response latency of trigeminal afferents during application of the pure odorant phenylethyl alcohol (PEA, 50% v/v) to the solution perfusing the bath (n=6, data not shown).

**Figure 4:**
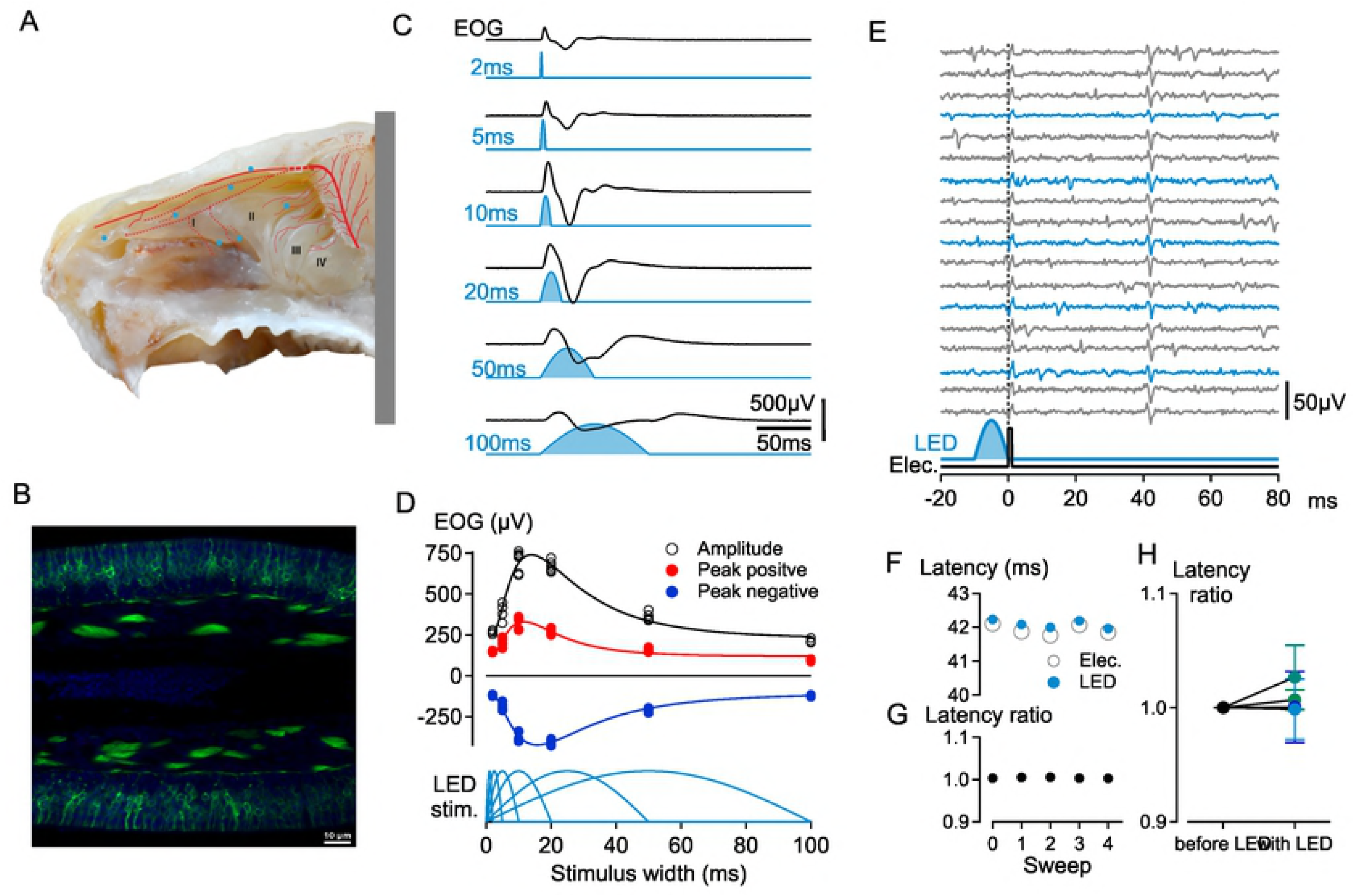
Optogenetic assessment of functional interactions between olfactory and trigeminal sensory afferents in the nasal cavity. Electrical receptive field locations for seven trigeminal afferents (A, blue markers) recorded in OMP/ChR2-YFP mice. Panel B shows the fusion protein ChR2-YFP (green) and nuclear DAPI (blue) staining in the olfactory epithelium in transverse section. Photoactivation of olfactory sensory neurons was verified by recording extracellular eletro-olfactogram (EOG) signals with an electrode positioned on the second turbinate (A, II) in response to sinusoidal light pulses varying in duration from 2-100ms (C). The absolute amplitude of the EOG signal was maximal in response to a 14 ms sinusoidal light pulse (D, black trace), while the positive going EOG was maximal for stimuli of 9 ms duration (D; red trace) and the negative-going EOG signal had a maximum amplitude at stimulus widths of 18 ms (D; blue trace). The response latency of action potentials in single trigeminal axons evoked by electrical stimulation (E, lower black trace) was monitored during electrical stimulation alone every 4s (E, grey traces) and combined with photoactivation (E, lower blue trace) of olfactory sensory neurons every 12s (E, blue traces). The average response latency to electrical stimulation alone (F, open grey markers) was compared to the electrical response latency when applied with light stimulation (F, blue markers) and the ratio of these two latencies determined (G). Pooled latency ratios for electrically-evoked responses in trigeminal afferents without light stimulation (control) and in combination with photostimulation (LED) are shown for seven fibres in panel H.

### Functional assessment of collateral branching in trigeminal sensory axons to the nasal cavity and olfactory dura mater

A functional pathway of communication between the olfactory epithelium and the olfactory bulb has been proposed (Schaefer et al, 2002) in which axon reflex conduction of action potentials allows signaling between sibling branches of individual trigeminal axons in the ethmoid nerve. To examine this interesting idea, we took advantage of the ability to record from spatially distinct sites along the ethmoid nerve (Figure 5) We began by recording from the distal cut end of the ethmoid nerve cut immediately distal to its entry point into the anterior cranial fossa through the ethmoid foramen (Figure 5B&E). Recording at this site, it was possible to find receptive fields of single C-fibres using an electrical stimulus in either the olfactory dura mater (Figure 5A) or the nasal cavity (Figure 5C). Afferents responses to mechanical stimulation were also observed using either a von Frey hair applied manually to sites in the olfactory dura (Figure 5D) and nasal cavity (Figure 5F) or a servo-driven mechano-stimulator in the olfactory dura mater or the nasal cavity (data not shown). This was entirely consistent with the topography of ethmoid innervation revealed by anterograde tracing (Figure 2A) and established that the ethmoid nerve provides functional innervation to structures both within nasal cavity and within the anterior carinal fossa, specifically the cranial meninges.

**Figure 5:**
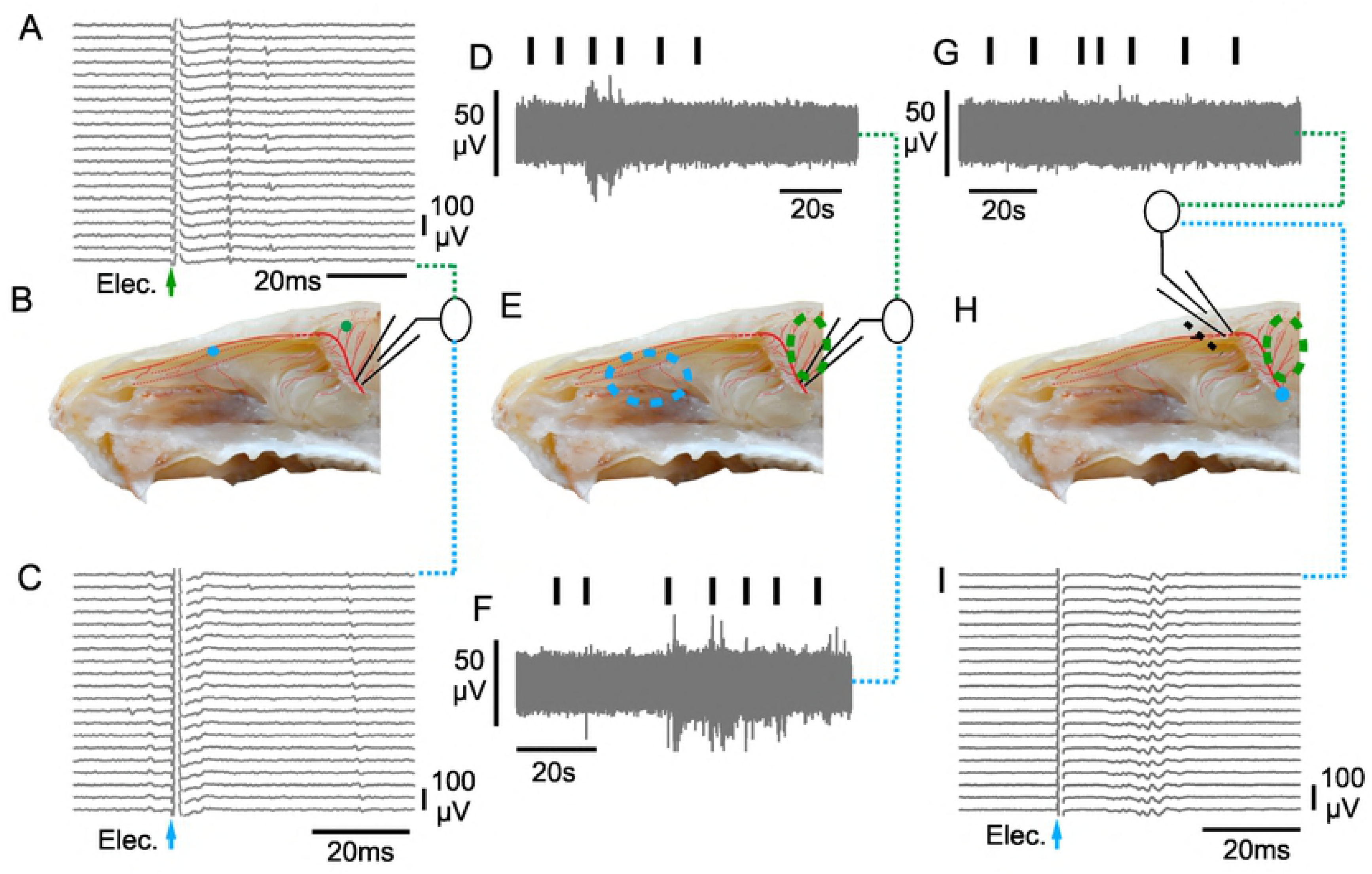
Functional assessment of potential axonal branching of individual trigeminal afferents to the olfactory epithelium and the olfactory dura mater. Recording from the distal cut end of the ethmoid nerve at its entry into the anterior cranial fossa (B) it was possible to verify electrical (A&C) receptive fields for single trigeminal axons both in the nasal cavity (C) and in the olfactory dura mater (A). Using the same recording configuration, it was possible to verify action potential activity (D & F) in response to mechanical von Frey probing (D & F, vertical black markers) in the nasal cavity (E, dashed green circle) and in the olfactory dura mater (E, dashed blue circle). In the same half-skull preparation as shown in panels D through F, the position of the recording electrode was changed to a more distal site, specifically the proximal cut end of the anterior ethmoid nerve after cutting it upon its entry into the nasal cavity through the cribroethmoid foramen (H). At this site, it is only possible principally to record anti-dromic activity in the trigeminal axons. We verified this using electrical stimulation to evoke a multi-fibre compound C-fibre action potential response (I) when stimulating at the original site of recording on the ethmoid nerve (H, blue dot). Functional assessment of whether axon reflex signals could propagate from sites in the anterior cranial fossa anti-dromically into the nasal cavity by stimulating with a von Frey filament (G; black vertical markers) at sites within the olfactory dura mater (G, dashed green circle). Mechanically-evoked activity was not observed (G).

To test formally whether this macroscopic branching of the ethmoid nerve comprised individual trigeminal axons with divergent branches innervating both olfactory dura mater and nasal structures we recorded from the proximal cut end of the ethmoid nerve cut immediately distal to its entry point into the nasal cavity through the cribroethmoid foramen (Figure 5H). Any action potential activity recorded from the electrode positioned at this site must be travelling in an anti-dromic fashion, i.e. towards sensory nerve terminals within the nasal cavity. To confirm that it was possible to record action potential activity in this manner, the stimulation electrode was placed on the parent ethmoid nerve at a site immediately distal to the ethmoid foramen (Figure 5H, blue dot) and a time-locked compound C-fibre action potential, i.e. multi-unit response, was evoked using constant current pulses (Figure 5I). We then searched for functional evidence of branching in individual axons by probing the olfactory dura mater with a von Frey probe. If individual axons branched to both the olfactory meninges/bulb and the nasal cavity, axon reflex conducted action potentials should have been evident at the recording electrode in the nasal cavity in response to mechanical probing of the olfactory dura mater. No activity was seen in this preparation (Figure 5G) nor were we were able to discern evoked action potential activity in any of six further preparations. This suggests that individual axons rarely branch to innervate both the olfactory epithelium and olfactory dura mater. In which case, it is difficult to consider axon reflex signaling a distinct functional pathway able to mediate cross-talk between trigeminal signals arising in the olfactory epithelium and neuropeptide release in the olfactory dura mater.

## Discussion

This study set out to examine interactions between the dual olfactory and trigeminal chemosensory systems in the nose. Using a forced choice behavioural assay, we found that a combination of odorant and irritant stimuli mitigated the otherwise prominent aversion of mice to irritant stimuli alone. Previous reports suggest that cross-talk between olfactory and trigeminal chemosensory signals might take place within the nose, either through paracrine effects mediated by local release of neurotransmitters [24] or through axon reflex signaling in branched trigeminal afferents (Schaefer et al 2003). We tested each of these proposals by recording directly from trigeminal axons innervating the nasal cavity. Using optogenetic techniques to activate exclusively olfactory sensory neurons we were unable to verify any modulation within individual trigeminal afferent axons. We also found no evidence for axon reflex signaling within individual trigeminal axons in the ethmoid nerve that might otherwise form a nexus between the olfactory epithelium and the olfactory bulb. An inability to verify cross-modal interactions between olfactory and trigeminal structures within the nasal cavity suggests that behavioural manifestations of olfactory-trigeminal cross-talk are most likely to occur at more central sites such as the trigeminal brainstem nuclei.

Functional recordings from trigeminal axons innervating the nasal cavity have shown that afferent signals are generated in response to a range of stimuli including odorant and irritant chemicals [50-52], mechanical probing in the nostrils [53] and cooling [54]. We verify here the ability of trigeminal afferents to encode each of these sensory modalities and extend the range of stimuli to include sensitivity to heat. In addition, we found that akin to somatosensory afferents in skin [55] and trigeminal ganglion neuronal somata [56] many trigeminal afferents in the nasal cavity are polymodal with heat thresholds around 45°C and mechanical activation thresholds similar to those reported for individual meningeal trigeminal afferents [40, 57]. Notably, we saw no evidence of warm fibres but did observe occasional cold-sensitive units as part of the background activity during heating and cooling ramps. The olfactory epithelium receives a lower density of trigeminal afferents than the respiratory epithelium [6]. Our functional mapping of trigeminal afferents in the nose confirmed the lower density in the olfactory epithelium (Figure 3). Using viral tracing methods to select somata from trigeminal ganglion neurons innervating the nasal cavity, a higher incidence of chemosensitive neurons in particular neurons expressing were encountered [58]. We confirm here that trigeminal sensory axons within the nasal cavity were often chemosensitive but in our hands units terminating within the olfactory epithelium were not activated by mechanical stimuli (Figure 3).

Human psychophysics indicate that odorants can act as irritants and likewise most irritants have an odour [15]. However, the threshold for chemesthetic trigeminal activation is typically an order of magnitude higher than that for olfactory sensory neurons [52]. Accordingly, combinations of odorant and irritant typically lead to predominance of the sensation of pungency, attributable to activation of trigeminal afferents, at the expense of odour perception [14]. While it is well established that odour perception [5], odour detection [19] and odour localization [59] are all affected by concomitant trigeminal activation, there is very little information as to whether olfactory stimulation might affect trigeminal activation. Recent observation in people using odour localization as an index of trigeminal activity, indicate that the presence of odorants can enhance trigeminal perception and attributed this to an interaction within the nose [26]. Our findings here using an isolated preparation of the mouse nose suggests that paracrine effects on trigeminal chemosensory signaling of olfactory sensory neuron activation are not apparent at the level of action potential generation (Figure 4). Although the use of photoactivation of olfactory sensory neurons excludes non-specific actions associated with chemical application, we cannot rule out the possibility of volatile stimuli applied to humans or mice affecting other chemosensory cells. For example TRPM5-expressing solitary chemosensory cells are located in the main olfactory epithelium of rodents [60] and are potentially capable of vesicular release of humoural mediators [61] which in turn could act in a paracrine manner on trigeminal nerve terminals.

In addition to paracrine interactions within the nasal cavity, Schaefer et al (2002) have suggested that the branching of individual trigeminal sensory axons to innervate both the nasal cavity and the olfactory bulb may constitute a pathway subserving trigeminal-olfactory interactions. In this scheme, a branched axon may constitute a signaling pathway through axon reflex conduction of action potentials from a site of generation in the terminals of one branch and subsequently antidromically to sister branches to effect neurotransmitter release. This idea is well established in cutaneous sensory nerve terminals where axon reflex mediated release of vasoactive neuropeptides gives rise to the spreading flare around a site of injury [62]. Some functional reports also suggest that proximal axonal branching in individual sensory axons enables innervation of spatially separate tissues [63]. Consistent with this concept, structural studies indicate that trigeminal nerves branch divergently to innervate meninges, cranial bone and extracranial periosteum [64]. Subsequent functional studies using collision techniques confirmed branching individual sensory axons in the spinosus nerve that gave rise to discrete mechanical and electrical receptive fields in intracranial meningeal structures and in extracranial muscle and fascia [65]. This observation extends the concept that headaches arise from activation of trigeminal afferents innervating the cranial dura mater to one in which the origin of headache includes activation of their axonal projections at extracranial sites. Similarly, irritant chemical stimuli could potentially trigger headaches [66] by activating trigeminal afferent innervating the nasal cavity with collateral branches to the olfactory dura mater. Therefore we adopted similar functional techniques to those of Schueler, Messlinger (65) but were not able to confirm the observation by Schaefer, Bottger (42) for axons in the ethmoid nerve, at least not for individual axons that might branch to innervate the nasal cavity and the olfactory dura mater (Figure 5). Although it is not possible to assess all axons within the ethmoid nerve, the absence of any retrograde axon reflex action potential activity suggests that if branching between these sites does occur, the incidence is likely to be rather low.

Human perception of odours can be modified by the presence of irritants. Similarly, we observed that irritant aversion in mice can be mitigated by co-application of an odorant. Both observations imply an interaction between chemical activation of nasal trigeminal and olfactory pathways at a level sufficient to affect behavior. On the basis of direct recordings from trigeminal sensory axons innervating the nasal cavity, it is not likely that this behavioural effect is causally related to interactions within the nose, implicating the trigeminal brainstem nuclei or higher convergent brain areas as sites for sensory cross-talk between trigeminal olfactory chemosensory signaling.

## Acknowledgments

We are grateful to Anja Bistron, Birgit Vogler and Gabi Guenther for their reliable and proficient technical assistance. This work was supported by a Deutsche Forschungsgemeinschaft (DFG) grant SFB1158-Project A04 to RWC and SF and FP7 project EUROHEADPAIN (Grant agreement no: 602633) to KM.

